# Nematode species differ in susceptibility and response to the nematode-trapping fungus *Arthrobotrys oligospora*

**DOI:** 10.64898/2026.11.04.764212

**Authors:** Dominika Lastik, Mara Carey-Wood, Rhiannon Wright, Henry Skitt, Thomas Sinanan, Freya Cole, Aine Murphy, Rebecca Forey, Daniel Henk, Vicky L Hunt

## Abstract

Fungal predators of nematodes are widespread and represent promising candidates for biological control of parasitic nematodes. However, the behavioural and molecular mechanisms underlying nematode–fungus interactions remain poorly understood, especially for nematodes other than *Caenorhabditis elegans*. We investigated the interactions between the nematode-trapping fungus *Arthrobotrys oligospora* and three nematode species: (i) free-living *C. elegans*, (ii) fig-associated *C. inopinata*, and (iii) the free-living adult life cycle stage of the gastrointestinal parasite *Strongyloides ratti*. *A. oligospora* trapped *S. ratti* more efficiently than either *C. elegans* or *C. inopinata*. Behavioural analyses revealed species-specific differences in nematode locomotion. *S. ratti* moved slower, travelled shorter distances and showed lower displacement than *Caenorhabditis* spp. To test whether locomotion influences fungal trapping, we examined eight *C. elegans* mutants with hyper- or hypoactive locomotory phenotypes. Although both groups moved at similar speeds, hyperactive mutants showed greater displacement, which positively correlated with trap formation but did not explain trapping efficiency by *A. oligospora*. We investigated the fungal response during interaction with the three nematodes. Exposure to *S. ratti*, but not to *Caenorhabditis* spp, resulted in reduced sporulation and differential expression of sporulation-associated genes. In nematodes, myosin-related genes responded to *A. oligospora* in a species-specific manner, including upregulation in *C. elegans* and downregulation in *S. ratti*. These transcriptional changes may reflect alterations in muscle structure, including reduction in dense body and mitochondrial size in *Caenorhabditis* species but not *S. ratti* during *A. oligospora* exposure. Together, our findings demonstrate that both fungal and nematode responses are nematode species dependent. These results improve our understanding of nematode–fungus interactions and highlight the importance of nematode species when developing nematode-trapping fungi as a biological control agent.

## 1. Introduction

Nematodes are the most abundant animals on earth [1] and inhabiting diverse environments all over the world [2]. Across these environments, nematodes interact with diverse organisms including nematophagous fungi which capture and pray on nematodes [3,4]. Nematophagous fungi have gained interest as a potential biological control agent of plant-parasitic [5,6] and animal-parasitic nematodes [7,8]. Based on their mechanisms of predation, nematophagous fungi can be broadly classified into three groups: 1) Nematode Trapping Fungi (NTF), 2) parasitic fungi and 3) toxin-producing fungi [9,10]. Here we focus on NTFs and their interactiosn with different nematode species. The most studied and the most commonly isolated NTF is *Arthrobotrys oligospora*, which, under starvation conditions develop spcialised loop-like trapping structures to capture nematodes, followed by digestion to obtain nutrients [11,12].

*Arthrobotrys* spp. encounter and trap a broad range of nematode species [13], and, *A. oligospora* can trap nematodes isolated from habitats distinct from its own, suggesting it is a generalist predator [14]. Less is known about how predation efficiency of *A. oligospora* varies between nematode species, although it is suggested to vary, with some nematodes such as *Ditylenchus destructor* observed to have higher survival rates [15,16]. Interestingly, a study comparing trapping efficiency rate of 26 different *Arthrobotrys* isolates against a free-living (*Panagrellus sp.*) and a parasitic nematode(*Cooperia punctata*) found that all isolates trapped fewer *C. punctata* nematodes [17]. These findings suggest that despite its broad prey range, the predation efficiency of Arthrobotrys may vary considerably amongst nematode species.

*Caenorhabditis elegans*, a free-living nematode, has been well documented for its interaction with *A. oligospora.* Although little detail is known about *A. oligospora* interactions with nematodes other than *C. elegans*. When *A. oligospora* grows with *Caenorhabditis* nematodes, it mimics food and pheromone cues to attract nematodes [18,19] that it then uses as food in low-nutrient environments [20,21]. After sensing it’s nematode prey, *A. oligospora* upregulates genes contributing to adhesion, fungal stress response, signalling, pH sensing, calcium binding, metabolism and transcriptional regulation leading to the formation of traps, as well as abundant secretion of proteins such a peptidases and carbohydrate binding protein domains [22–25]. Less is known about transcriptional changes in nematodes when exposed to NTF. *C. elegans* upregulate genes coding for galactose-binding lectins, protease inhibitors and neuropeptide proteins when exposed to the NTF *Dactylellina haptotyla*, [26] but to our knowledge, there is no published transcriptome data in nematodes exposed to *A. oligospora*.

Here, we investigated interactions between the nematode-trapping fungus *A. oligospora* and three nematode species: (i) free-living *C. elegans*, (ii) fig-associated *C. inopinata*, and (iii) the free-living adult life cycle stage of the gastrointestinal parasite *Strongyloides ratti*. We identified species-specific changes in gene expression in both *A. oligospora* and the nematodes, as well as physiological changes in the fungi - sporulation and growth - and in the nematode – muscle tissue.

## 3. Results

### 3.1 A. oligospora is more efficient at trapping S. ratti than Caenorhabditis spp

We investigated *A. oligospora* trap formation when cultured with a single nematode species, either *S. ratti, C. elegans* or *C. inopinata* (**Fig. 1A**). When cultured with *S. ratti*, *A. oligospora* produced 61.1 ± 35.4 (mean ± se) traps which is significantly fewer traps than the 444.3 ± 126.7 *A. oligospora* traps produced when incubated with *C. elegans* (Kruskal-Wallis multiple comparison test p-value < 0.0001). *A. oligospora* produced 214.5 ± 70.1 traps when exposed to *C. inopinata* (**Fig. 1B, SI Table 1**). For all three species, it was necessary for nematodes to be alive to induce *A. oligospora* trap formation indicating that the fungus detects the nematodes when they are alive (**SI Fig 1, SI Table 1**). Overall, *A. oligospora* was more efficient at catching *S. ratti* nematodes in traps and 83% of nematodes were trapped after 24 hours, even though fewer traps were produced overall. In contrast 37% and 27% of *C. elegans* and *C. inopinata* were trapped after 24 hours (**Fig. 1C, SI Table 1**).

**Figure 1.**
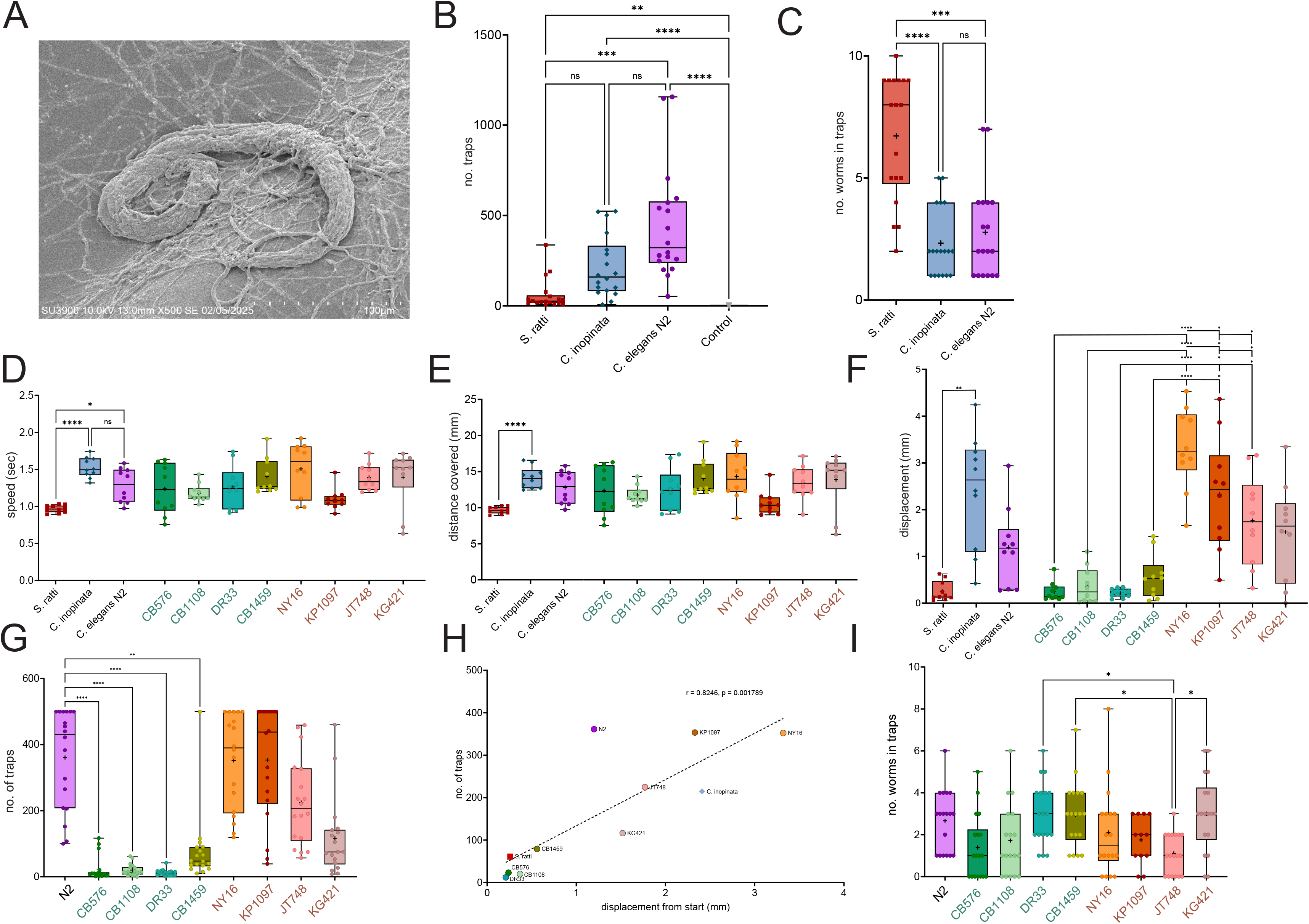
*A. oligospora* trap formation when exposed to nematodes. (A) SEM image of *S. ratti* nematode trapped by *A. oligospora.* (B) Number of traps formed on LNA plates after 24 hours of exposure to 10 *C. elegans* adults (mean 444.3 ± 126.7)*, C. inopinata* adults (214.5 ± 70.1) or *S. ratti* (61.1 ± 35.4) free-living adults (significance calculated using Kruskal-Wallis multiple comparison test. (C) Proportion of nematodes trapped after 24 hrs for *C. elegans* (37%)*, C. inopinata* (27%) and *S. ratti* (83%) (Kruskal-Wallis multiple comparison test). (D) Speed (sec) (E) distance travelled (mm) and (F) displacement (mm) of *C. elegans* lab strain (N2), four hypoactive *C. elegans* lines: CB576, CB1108, DR33 and CB1459, four hyperactive *C. elegans* lines: NY16, KP1097, JT748 and KG421, *C. inopinata* and *S. ratti*. (Brown-Forsythe and Welch ANOVA multiple comparison test, p<0.0001). (G) Number of traps formed and (H) correlation of number of traps vs displacement from start position (mm) for all nematodes. Statistical comparisons are located in SI Table 1) Pearson’s correlation coefficient (r) and p value calculated in GraphPad, Prism. (I) number of worms trapped after 24 hours exposure to 10 nematodes on LNA plates with *A. oligospora* in response to *C. elegans* N2, four hypoactive *C. elegans* lines, four hyperactive *C. elegans* lines, *C. inopinata* and *S. ratti*. (Kruskal-Wallis, p = 0.0002). Statistical analysis carried out GraphPad, Prism. Results considered significant if p value = * < 0.05, ** <0.01, *** <0.001, **** <0.0001.

Free-living adult *S. ratti* are a similar size to *C. elegans* adult females (∼1mm [27,28] but their locomotive behaviour differed from *C. elegans* and *C. inopinata. S. ratti* moved at a significantly slower speed than *C. elegans* and *C. inopinata*, over shorter distances and had lower displacement (i.e. stayed closer to the origin location) than *C. inopinata* on NGM plates (Brown-Forsythe and Welch ANOVA multiple comparison test, p value <0.0001) (**Fig. 1D-F**, **SI Table 1**). We hypothesised that the reduced speed and displacement of *S. ratti* make these nematodes easier for *A. oligospora* to locate and trap. To test this, we characterised the movement of four hypoactive (CB576, CB1108, DR33 and CB1459) and four hyperactive (NY16, KP1097, JT748 and KG421) *C. elegans* mutant lines (**Fig. 1D-F, SI Figure 2, SI Table 1**). The mutant lines all moved at a similar speed over a similar total distance. However, the hyperactive lines had a higher displacement value, in line with those observed for *C. elegans* N2 and *C. inopinata,* and the hypoactive lines had a lower displacement value, similar to *S. ratti* (**Fig. 1D-F**). The hyperactive lines induced more *A. oligospora* traps than the hypoactive lines (Kruskal-Wallis multiple comparison test p-value < 0.0001) **Fig. 1G)**. Overall, the displacement distance from the origin point was strongly correlated (r = 0.8246, p = 0.001789) with the number of traps formed **(Fig. 1I)** but no significant correlation (r = 0.18 and 0.27, p = 0.5865 and 0.41) was observed between number of traps formed and either total distance (mm) or speed (sec) (**SI Figure 3**). The number of nematodes trapped was the same across the hypoactive and hyperactive lines indicating that displacement value alone is not associated with trapping success and other factors are responsible for species-specific trapping (**Fig. 1H, SI Table 1)**.

**Figure 2.**
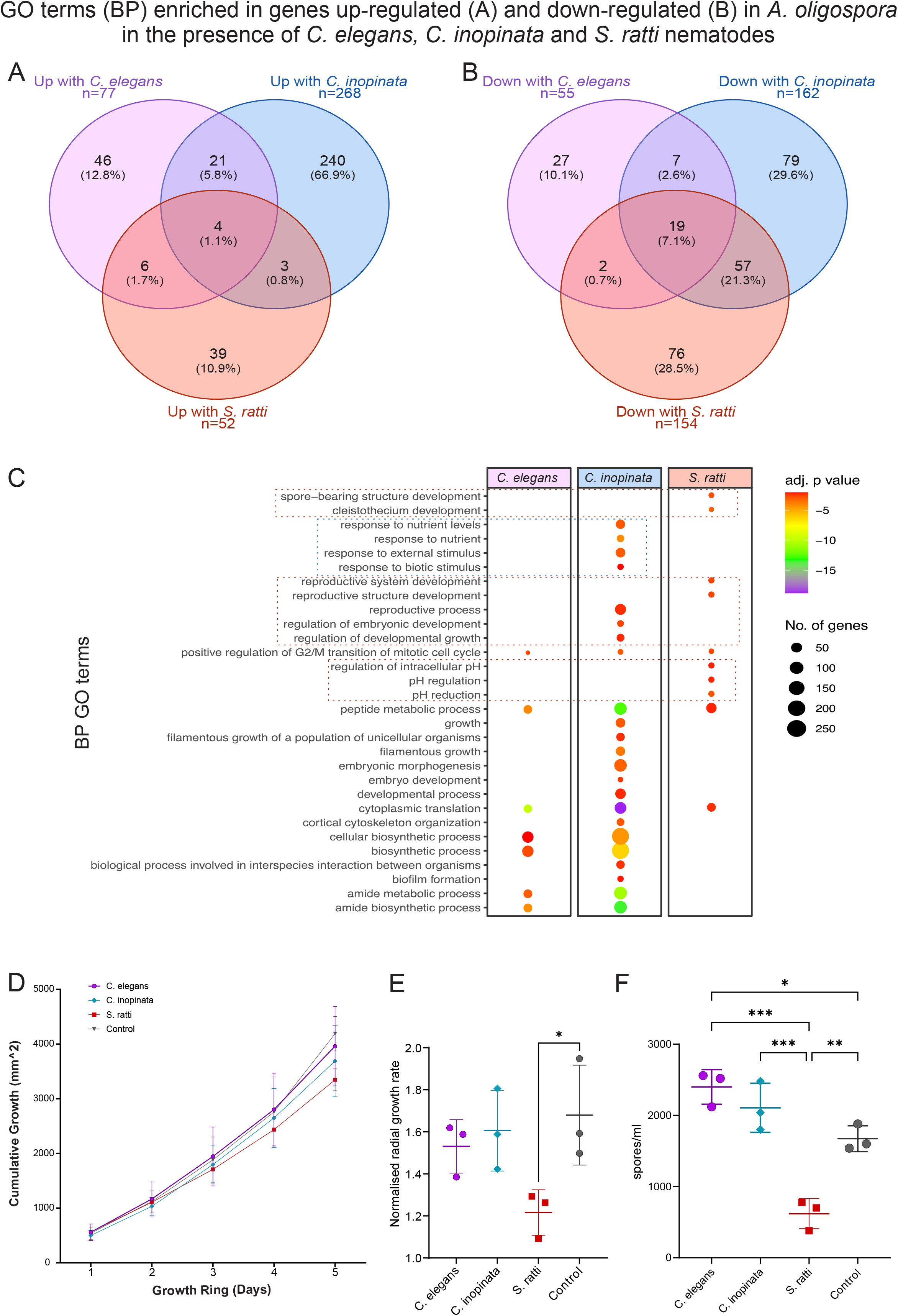
Genes associated with sporulation are uniquely upregulated in *A. oligospora* when exposed to *S. ratti*. Gene Ontology (GO) terms enriched in (A) upregulated and (B) downregulated genes in *A. oligospora* incubated with *C. elegans*, *C. inopinata* and *S. ratti* nematodes compared to *A. oligospora* incubated without nematodes (DESeq2 adj. p value ≤ 0.01 and log fold change (FC) > 1). (C) Biological Processes GO terms upregulated in *C. elegans*, *C. inopinata* and *S. ratti* involved in embryonic development and sporulation. (D) Cumulative growth (area) of *A. oligospora* in the presence of nematodes cf. no nematode control across a five-day incubation. (E) Radia growth rate of *A. oligospora* in the presence of nematodes vs no nematode control in five days on LNA. (F) number of spores produced by *A. oligospora* in response to incubation with and without nematodes on LNA (one-way ANOVA with Tukey’s multiple comparisons test). Results considered significant if p value = * < 0.05, ** <0.01, *** <0.001, **** <0.0001.

**Figure 3.**
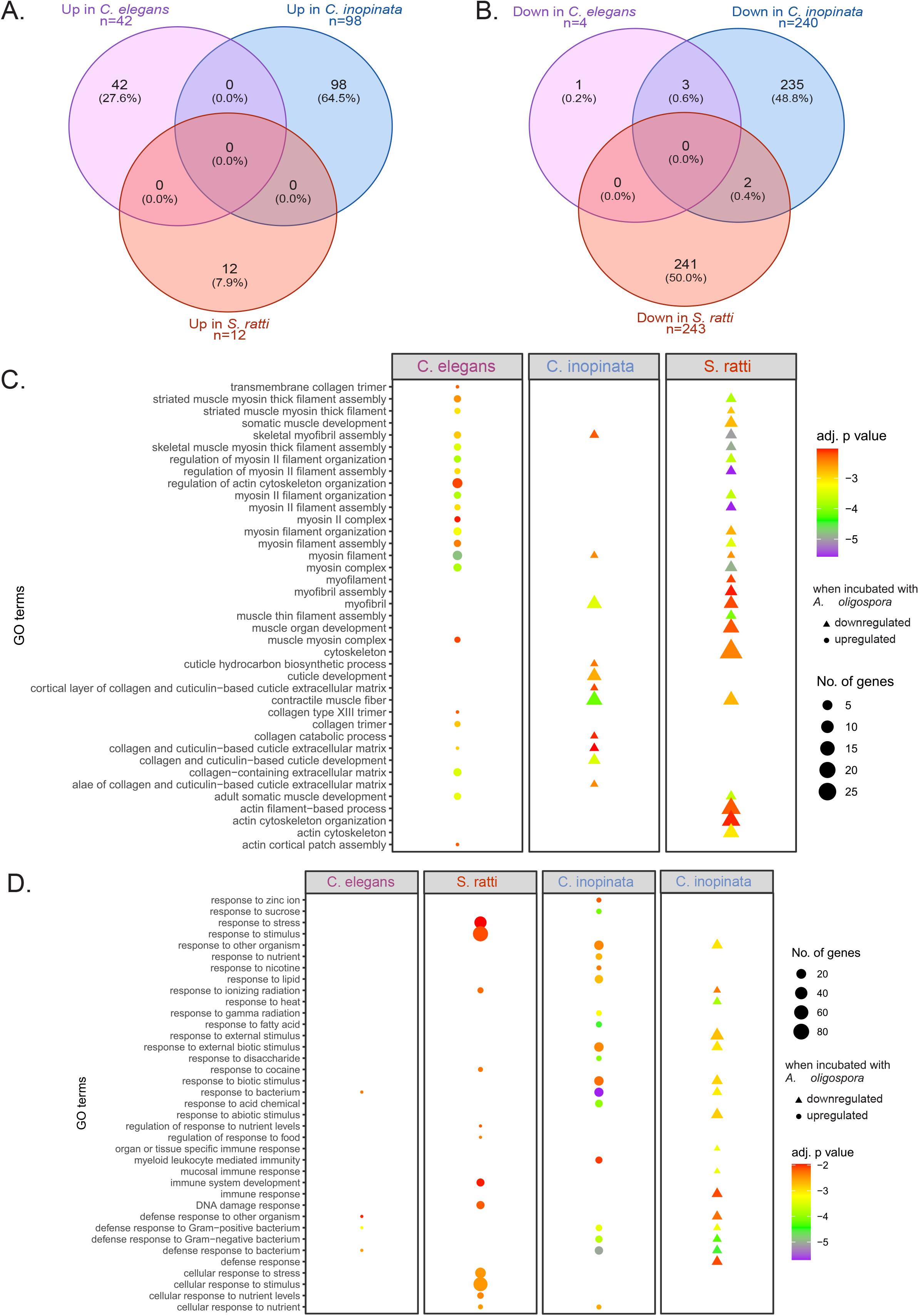
Myosin-related genes upregulated in *C. elegans* but downregulated in *S. ratti* and *C. inopinata*. Gene Ontology (GO) terms enriched in (a) upregulated and (b) downregulated genes in *C. elegans*, *C. inopinata* and *S. ratti* when incubated with *A. oligospora* compared to nematodes not incubated with *A. oligospora* (DESeq2 adj. p value ≤ 0.01 and log fold change (FC). GO terms related to (c) muscle, actin and myosin (d) and defence or response mechanisms enriched from differentially expressed genes in *C. elegans*, *C. inopinata* and *S. ratti* when incubated with *A. oligospora*. Colour indicated by adjusted p value, size indicated by number of significant genes. GO terms reduced with Revigo and graph visualised in R (version 4.2.1).

### 3.2 *A. oligospora* upregulates pH regulation- and sporulation-associated genes and exhibits reduced sporulation in response to *S. ratti*, but not *Caenorhabditis* spp

To better understand the mechanisms underlying differences in trap induction and trapping efficiency among nematode species, we characterised the transcriptional response of *A. oligospora* following exposure to *S. ratti*, *C. elegans*, or *C. inopinata*, and compared these responses with fungus grown in the absence of nematodes. Exposure to *S. ratti* resulted in the differential expression of 2,054 genes (1,150 upregulated and 904 downregulated), whereas exposure to *C. elegans* altered the expression of 828 genes (414 upregulated and 414 downregulated). The largest transcriptional response was observed following exposure to *C. inopinata*, with 2,931 differentially expressed genes (1,443 upregulated and 1,488 downregulated) (**SI Table 2A, SI Figure 4-5**). GO enrichment analysis of differentially expressed genes identified 19 downregulated Biological Process (BP) GO terms commonly associated with *A. oligospora* when exposed to all three nematode species (**Fig. 2A, SI Table 2B**). These GO terms included terms relating to fatty acid metabolic processes (GO:0006631, GO:0006633, GO:0042759), carboxylic acid biosynthesis (GO:0032787, GO:0019752, GO:0046394, GO:0072330) and sulphur compound metabolic process (GO:006790, GO:0044272, GO:0000096, GO:0000097). Genes associated with a further four BP GO terms were upregulated in *A. oligospora* when exposed to all three nematode species, including two GO terms associated with positive regulation of the G2/M stage of mitosis (GO:0010971, GO:1902751).

**Figure 4.**
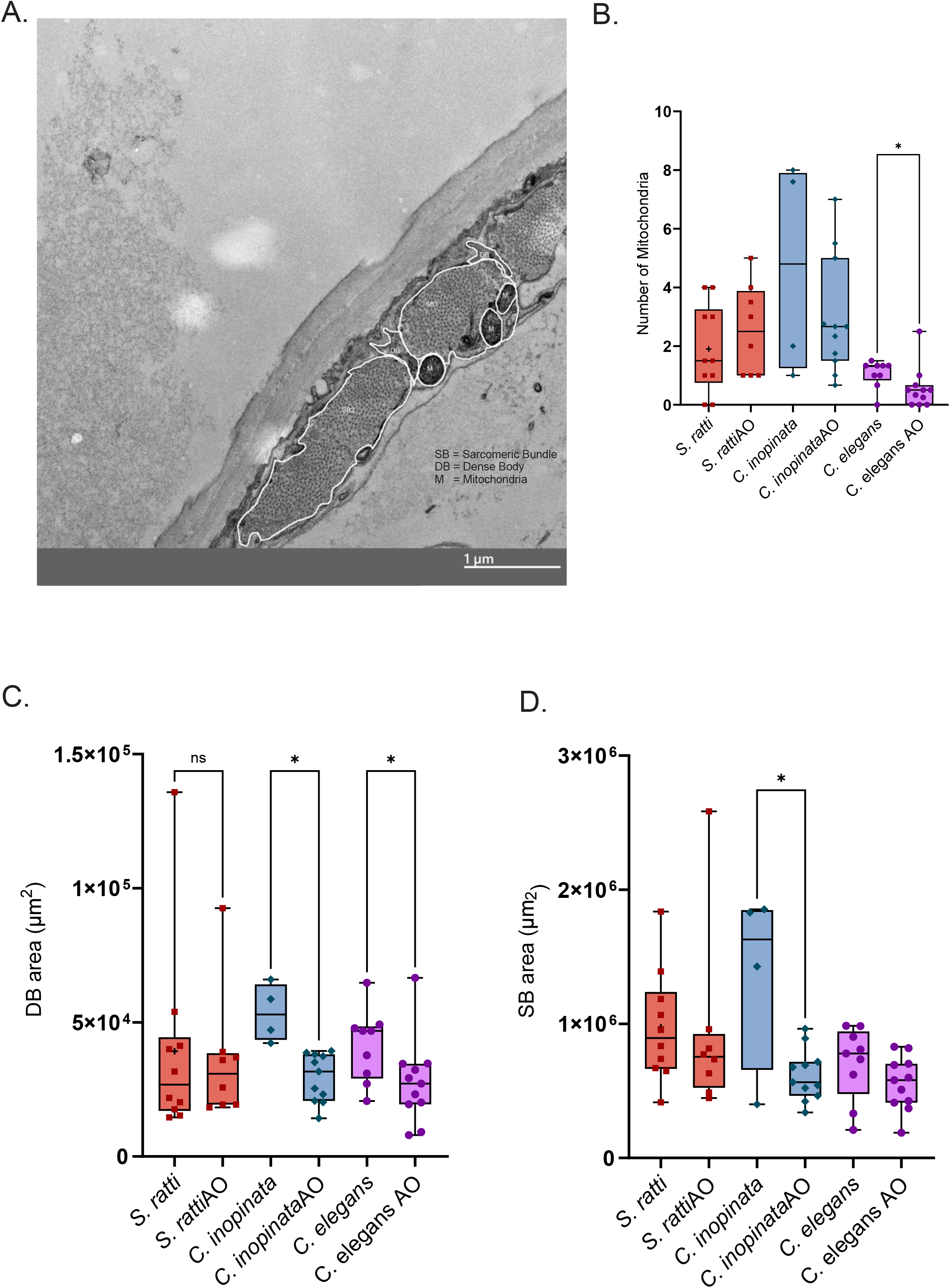
Cuticle and myosin TEM image analysis. (A) An example of an annotated *C. elegans* TEM cross section (11000x magnification) showing sarcomeric bundles (SB), dense bodies (DB) and mitochondria (M). (B) Number of mitochondria counted in nematodes vs nematodes incubated wit *A. oligospora* (Mann-Whitney’s test p = 0.0221). (C) Dense body (DB) and (d) sarcomeric bundle (SB) area of nematode muscle with and without exposure to *A. oligospora* (Kruskal-Wallis uncorrected Dunn’s test). Results considered significant if p value = * < 0.05, ** <0.01, *** <0.001, **** <0.0001.

The majority of enriched GO terms were associated with nematode species-specific changes. When cultured with *S. ratti*, 115 BP GO terms were uniquely associated with differentially expressed genes in *A. oligospora* including an upregulation of genes involved in pH regulation, particularly reduction in pH (GO:0030641, GO:0051452, GO:0051453, GO:0006885, GO:0045851), and spore production (GO:0000909, GO:0030582, GO:0030584, GO:0075259) (**Fig. 2B, SI Table 2b**). Interestingly, the response of *A. oligospora* to *C. inopinata* had more overlapping GO terms with *S. ratti* than its close relative *C. elegans,* including 57 GO terms associated with genes downregulated by *A. oligospora* in both *C. inopinata* and *S. ratti* (**SI Table 2b**). Of these, seven downregulated genes related to GO terms associated with sporocarp formation or spore bearing structures and nine were related to GO terms associated with reproductive structure development. Enrichment of these GO terms was not observed for *A. oligospora* when incubated with *C. elegans* (**Fig. 2C**).

To determine whether the transcriptomic changes were associated with phenotypic differences, we investigated the effects of nematode presence on the growth and sporulation of *A. oligospora*. We measured cumulative growth to determine total area of growth and radial growth rate to determine average distance grown from the centre of plate (**SI Fig 6, SI Table 3**). No significant difference was observed in cumulative growth rate of *A. oligospora* when co-incubated with nematodes (**Fig. 2D**). However, a significant decrease in radial growth was observed in *A. oligospora* incubated with *S. ratti cf.* no nematode controls (ANOVA with Tukey’s multiple comparison, p=0.0462) (**Fig 2E**). In the presence of *S. ratti*, *A. oligospora* produced significantly fewer spores (620/1 ml) than controls (1,673/1 mL), *C. elegans* (2,400/1 mL) and *C. inopinata* (2,107/1 mL) (p = <0.05, one-way ANOVA with Tukey’s multiple comparison test). The number of *A. oligospora* spores produced in the presence of *C. elegans* was significantly (p=0.0321) higher than control. (**Fig 2F**). Together the transcriptome and sporulation data indicate that *A. oligospora* sporulates less when *S. ratti* is present and more when *C. elegans* is present.

### 3.3 Species-specific regulation of myosin-related genes following exposure to *A. oligospora* is accompanied by muscle-associated changes in *C. elegans*

To gain further insight into species-specific interactions between nematodes and *Arthrobotrys oligospora*, we characterised the transcriptional responses of the nematodes following fungal exposure. The transcriptomic response was species dependent, and the magnitude of the transcriptional response varied among species, with substantial differences in the number of differentially expressed genes. When exposed to *A. oligospora,* 195 genes were downregulated and 91 genes upregulated in *S. ratti*, 10 genes downregulated and 84 genes upregulated in *C. elegans* and 552 genes downregulated and 408 genes upregulated in *C. inopinata* (DESeq2, FDR< 0.01, logFC > 1) (**SI Table 4**).

GO enrichment analysis indicated that the gene expression response in the nematode was largely species-specific. No enriched GO terms were shared between species for genes that were upregulated during exposure to *A. oligospora,* and a total of only 5 genes were shared between species for downregulated genes (**Fig. 3A-B**). Genes downregulated in *S. ratti* included terms of developmental and reproductive processes, response to stress and nutrients and myosin-related processes while terms downregulated in *C. inopinata* included collagen and cuticle development, response processes and biosynthesis regulation (**3C-D, SI Table 4, SI Figure 7**). GO terms associated with myosin were upregulated in *C. elegans* when exposed to *A. oligospora*, but downregulated in *S. ratti* under the same conditions, compared with controls (**Fig. 3C**). This suggests that myosin is important in the response to fungal predators, but the nematode response is species-specific. Myosin, a motor protein, drives muscle contraction [29,30] and could therefore be associated with increased or decreased movement when exposed to *A. oligospora*.

We reasoned that changes in myosin-related genes may be reflected in muscle physiology. We measured different metrics associated with muscle tissue including number of sacromeric bundles, sacromeric bundle size, dense body number, dense body size, and the number of mitochondria within proximity of muscle tissue (**Fig 4A**). We observed differences between all three control nematodes indicating fundamental differences between these nematodes. When exposed to *A. oligospora* the number of mitochondria within 1 µM distance of muscle tissue was significantly lower in *C. elegans* but the same in *C. inopinata* and *S. ratti* (unpaired Mann Whitney test p*=*0.0221). Dense body area was also significantly reduced in *C. elegans* and *C. inopinata* when exposed to the fungus but not in *S. ratti* dense bodies (p*=*0.0281 and 0.0159). Further, there is a decrease (p=0.0440) in SB area in *C. inopinata* when exposed to fungus. No other significant differences were observed in the muscle tissue (**Fig. 4B-D, SI Table 5**). We also observed species-specific changes in five GO terms associated with cuticle or collagen downregulated in *C. elegans* incubated with *A. oligospora* and nine GO terms upregulated in *C. inopinata* incubated with *A. oligospora* (**Fig 3C, SI Table 4**). While we do not find cuticle-related GO terms in *S. ratti*, we do find five and three genes with protein descriptions of ‘nematode cuticle collagen’ and ‘collagen alpha-chain’ in genes up or down regulated in *S. ratti* when in presence of *A. oligospora.* We analysed features of the nematode cuticle on SEM images, but no physiological changes were observed when nematodes were exposed to *A. oligospora* (**SI Table 5**).

## 4. Discussion

After 24 h exposure, *C. elegans* induced greater trap formation by *A. oligospora* than *S. ratti*, and *S. ratti* was trapped more efficiently. *S. ratti* showed significantly lower speed, distance travelled and displacement than *C. elegans*, leading us to hypothesise that nematode movement influences trap induction. Displacement, which captures the area covered a nematode, was strongly correlated with the number of traps formed by *A. oligospora* but not the ability for the fungus to trap nematodes. The increased trap formation when nematodes travel over larger distances could reflect the increased area of secreted nematode ascarosides, which *A. oligospora* uses to detect and subsequently induce traps against *C. elegans* [31]. We could not rule out if the slower speed of *S. ratti* made the nematodes more vulnerable to trapping by *A. oligospora*, and further investigation is required to understand if other locomotory characteristics are important for nematodes to escape trapping by *A. oligospora*. It’s also important to note that nematode locomotion was recorded in the absence of the fungus and we do not know if the locomotory behaviour changes during fungal exposure. Overall, the results suggest that *S. ratti* is more easily trapped by *A. oligospora,* and that *A. oligospora* is detecting *Caenorhabditis* species, but less able to trap them.

Molecular differences could also be responsible for nematode species-specific differences in trapping ability, for example, if *A. oligospora* can more easily detect a particular species. In *C. elegans* ascaroside pheromones trigger NTF trap development [31]. Ascaroside signalling is generally conserved among nematodes [32] and has roles in dauer larval development in *C. elegans*, and larval development in parasitic nematodes [33,34]. However, the structure and composition of these metabolites vary between parasitic species including *Strongyloides* [34] which could account for differences in *A. oligospora* trapping response. Other metabolites, such as ammonia from urea, function as a signal molecule in the fungus to initiate the lifestyle switch to form traps [35] and could also influence trap efficiency. *A. oligospora* also uses proteins on the fungal surface to interact with sugar molecules on the nematode cuticle, [36,37], which may be different between the free-living *C. elegans* and parasitic *S. ratti* and could potentially explain the alternate effects on fungal hyphal growth.

*A. oligospora* produces fewer spores in response to *S. ratti* than other nematodes. Furthermore, in *A. oligospora* incubated with *S. ratti* we find genes significantly enriched for GO terms related to sporulation, pH regulation and reproductive development. pH regulation is essential for trap formation [38,39] and also important in spore and fruiting body production and reproduction in fungi [38,39] and these upregulated genes could therefore contribute to regulating *A. oligospora* trap formation in response to *S. ratti.* Fungi often sporulate as an active survival and dispersal mechanism and can negatively correlate with nutrient acquisition [40–44]. Differences in sporulation may therefore directly reflect the increased nutritional content when trapped with *S. ratti*, because substantially more *S. ratti* nematodes are captured by *A. oligospora* compared with the other two nematode species.

We investigated differences in nematode gene expression when exposed and not exposed to *A. oligospora* to further explore nematode-species specific differences in *A. oligospora* interactions. Interestingly, when incubated with *A. oligospora*, *C. elegans* upregulate and *S. ratti* and *C. inopinata* downregulate genes enriched for GO terms related to myosin, cytoskeleton, muscle, collagen and cuticle. While no physical differences were observed in the nematode cuticle during *A. oligospora* exposure, a reduction in number of mitochondria near muscle, and the area of the dense body and sacromeric bundle in *C. elegans* and *C. inopinata* muscle was observed during exposure to *A. oligospora*. In *C. elegans*, changes in DB and SB structure organisation through mutation of localised proteins result in changes in locomotion and muscle strength [45] and more rapid locomotion is associated with shorter sarcomeres and region-dependant changes in sarcomere numbers [46]. The changes we observe in nematode muscle tissue during exposure to *A. oligospora* could therefore reflect increased locomotion e.g. to escape fungal traps. Our analysis of TEM images of muscle and cuticle measured the size of different structures but didn’t measure other structural changes that may occur. A previous study reported that a mutation in a *C. elegans nhr-66* which regulates expression of collagen in *C. elegans* cuticle, reduces the trapping ability of *A. oligospora* by 40%, even though no physical changes were observed in the cuticle [47].

Future work focusing on more detailed composition of cuticle and muscle could identify further changes that occur in response to fungal exposure, corresponding to the strong species-specific changes observed gene in differential expression of muscle and cuticle related genes.

Reduced expression of myosin and muscle development associated genes in *S. ratti* could also reflect efforts to directly repair damage caused by fungal trapping or could reflect higher proportion of trapped *S. ratti* nematodes compared to other species in the samples.

Overall, these findings highlight the importance of species-specific differences in interactions between nematodes and NFTs. This is particularly relevant for parasitic nematodes, which remain comparatively understudied in the context of fungal predation. The efficient trapping of *S. ratti* suggests that *A. oligospora* has potential as a biocontrol agent against the free-living stages of *Strongyloides* spp. Understanding the mechanisms underlying species-specific differences in trap induction and susceptibility will therefore be important for assessing the feasibility and optimisation of NTF as a targeted biocontrol strategy.

## 4. Materials and Methods

### Nematode cultures

*S. ratti:* strain ED321 was maintained in 4–6 week-old female Wistar rats by subcutaneous injection of 550 third stage infective larvae (iL3) obtained by faecal culture as described in Hunt et al., 2016. This work was performed under the authority of licences issued by the Animals (Scientific Procedures) Act 1986. Mixed sex adult worms are collected from faecal cultures 3 days post collection.

*C. elegans* (strain N2) and *C. inopinata* (strain NK35) were maintained on Nematode Growth Media (NGM) agar with Streptomycin (100 mg/ml) and fed with streptomycin-resistant *E. coli* OP50 and kept at 20°C. Mixed sex adult or young adult stages were collected using liquid NGM media for experiments and washed with ddH_2_O three times.

*C. elegans* mutants were ordered from Caenorhabditis Genetics Centre (CGC).

### Induction of nematode death

The dead nematodes which were used within an experiment were collected and washed before being snap frozen in liquid nitrogen and stored at -70°C. Prior to use, nematodes were thawed, counted and death established by absence to respond to touch, as described by [48]

### Fungal culture

*A. oligospora* strain ATCC 24927 (from Westerdijk Institute, KNAW) maintained in glass-fronted incubator with seasonal daylight cycles at 24°C on potato dextrose agar (PDA) or corn-meal agar (CMA) for spore collection and on low-nutrient agar (LNA) or in low-nutrient media (LNM) for experiments [49]

### RNA extraction and sequencing

5 spores per µL were added to LNM and incubated by gentle shaking at 24°C for 5 days. Either a) 200 young adult or adult mixed sex nematodes (*C. elegans, C. inopinata* or *S. ratti*) were added in 50 µl LNM and left to incubate for further 6 hours, visually confirming trap formation or b) 50 µl LNA was added without worms, as a control and c) 200 young adult or adults mixed sex nematodes (*C. elegans, C. inopinata* or *S. ratti*) in LNM were left for 6 hours in the same conditions. Three replicates were collected for each condition.

Biomasher II tubes (Nippi) were used in combination with a standard handheld homogeniser. The samples were homogenised prior to addition of Trizol with three cycles of dry ice incubation and homogenisation. For RNA extraction the standard Trizol protocol (Invitrogen) was amended with the following alterations: 1-Bromo-3-chloropropane (BCP) used instead of chloroform, RNA was precipitated with glycogen for 1-2h at -20°C, pellet was washed three times using 75% ethanol. Total RNA was quantified using a Nanodrop (DeNovix) and Bioanalyzer (Agilent Technologies). Only samples with a minimum RIN value of 4 were sent for RNA sequencing. Data available on Sequence Read Archive (SRA) (<u>PRJNA1537376</u>).

### Induction of Trap Formation

From CMA plates, fungal agar plugs were sub-cultured as described previously onto low-nutrient agar (LNA) [49] for 4 days and again for 3 days before use. To induce trap formation 10 alive or dead nematodes were added onto plates and incubated in at 24°C for 24 hours. All plates were stained with Lactophenol Cotton Blue (Pro-Lab Diagnostics) and washed with PBS and 0.1% Tween three times and again three times with PBS to fix and stain the fungus for trap visualisation and counting. Traps and nematodes trapped were manually counted using a stereo microscope (Meiji Techno). At least 18 replicates were obtained for each nematode condition.

### Fungal growth assays

5 mm fungal agar plugs from previously inoculated low-nutrient agar (LNA) plates [49] were transferred to new LNA plates. Plates were incubated at 24°C for 3 days. The agar plugs were then removed and 0, 10 or 50 nematodes were added in 100 µL ddH_2_O incubated in daylight at 24°C for 5 days. Fungal growth was measured every 24 hours by marking the end of hyphal growth on the plates. Plates were fixed and stained with 5 ml of lactophenolcotton blue (Pro-Lab Diagnostics) and incubated at room temperature for 5 minutes. Excess stain was washed trice in 0.1% PBSTween 80 (Sigma) solution, followed by three 0.1% PBS washes. Stained plates were left in cold room until imaging.

### Imaging of fungal growth

Pictures of whole plates were taken using a Nikon D3200 camera at 30 cm away, in a dark room. We took radius measurements bidirectionally and compared them to our automated area measurements using Spearman’s correlation. Radial and cumulative area growth quantification of plates was conducted using ImageJ (v. 1.54) [50]. Each image was calibrated using measuring tape. The image was converted to 8-bit and threshold adjusted using the OTSU method. Circumference and area of each 24-hour growth ring was measured in two directions – 0 and 90 degrees.

### Spore collection and counting

50 *C. elegans*, *C. inopinata* and *S. ratti* were added to LNA plates with agar plugs and left to grow for three weeks at 24°C. Spores were collected by adding 5 mL of 0.1% Tween and ddH_2_O suspension to the grown plates and agitating with a plate spreader, before collecting through a Miracloth filter. Spores were counted in 10 x 5 µL drops. Three replicate plates were counted per nematode species. All statistical analysis was conducted via GraphPad Prism.

### Statistical analysis of fungal growth and sporulation

Statistical analysis was conducted via GraphPad Prism. Initial relative growth rate (RGR) from each replicate was calculated using the following equation.

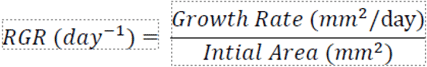

Growth rate (mm^2^/day) was calculated as the slope of the linear regression of cumulative area over time, normalised to the initial area obtained on day 1 of the experiments. Normalising the data allows for size-independent manner of growth allowing for comparisons between species, expressed in units of day [51,52]. Differences in the RGR of nematode species were analysed using the non-parametric Kruskal-Wallis test followed by Dunn’s multiple comparison test for post-hoc pairwise comparisons. To adjust for these multiple comparisons, Bonferroni’s correction was applied. Pairwise comparisons within species under light and dark conditions were performed using the Mann-Whitney U test. To adjust for these independent multiple comparisons, Bonferroni’s correction was applied to control for family-wise error rate. Adjusted significance was *p* < 0.0125.

### Measuring nematode cuticle

As in worms collected for RNA extraction, adult nematodes were added to a) *A. oligospora* growing in LNM or b) LNM only in a shaking incubator at 24°C for 5 days. *C. elegans, C. inopinata* or *S. ratti* nematodes were added in 50 µl LNM and left to incubate for further 6 hours, visually confirming trap formation. Three replicates were incubated for each condition. The nematodes were extracted, placed in 1 ml of fixative (glutaraldehyde + paraformaldehyde in sodium cacodylate buffer) and cut open diagonally in the middle using a scalpel and placed on ice. The samples were then processed for TEM imaging at the Wolfson Bioimaging Facility, University of Bristol.

To calculate the thickness of the nematode cuticle, ImageJ was used to lay a grid over the images to measure the cuticle at eight sections for consistent technical replicates for three biological replicates of all conditions. The area and perimeter were calculated by measuring the circumference of the nematode and the area of the cuticle was calculated by measuring the area of the nematode from the inner edge of the cuticle and subtracting this value from the total area of the nematodes. The thickness of each cuticle layer (surface coat layer, epicuticle layer, medial layer, fibrous layer and basal layer) was measured using ImageJ. Five technical replicates from different images were measured using a layover grid to ensure continuity between images.

Statistical analysis was preformed using GraphPad Prism. Unpaired t-test with Welch’s correction was used to compare the number of cuticle annuli, cuticle area, perimeter, total cuticle thickness and individual cuticle layer thickness between nematodes incubated with and without *A. oligospora.* One-way ANOVA was used to analyse differences in cuticle layer thickness amongst the uninfected nematode species. Results with p value <0.05 were considered significant. Effect sizes between samples were calculated using Hedges’ g, with 95% confidence intervals (CI). Effect sizes and CO were calculated using an online calculator (Calcarena, https://calcarena.com). Hedges’ g values effect sizes were classified as negligible (<0.20), small (0.20-0.50), moderate (0.50-0.80) and large (0.80+).

### Measuring nematode muscle changes

TEM images used to measure cuticle thickness were also used to determine differences in the area and morphology of the muscle. Three morphological structures were measured within these images, sarcomeric bundles (SB), dense body structures (DB) and mitochondria (M). Sarcomeric bundles are bundles of actomyosin fibres (both myosin and actin filaments) which make up the bulk of the muscle tissue. Dense bodies are a collection of actin filaments which divide the sarcomeric bundles. The number of complete sarcomeric bundles, dense bodies and mitochondria were counted. Mitochondria were only included in the count if they were within a 1 µm distance from the edge of a sarcomeric bundle. The area of all individual structures was measured, and a mean average was taken across all the complete structures within an image using ImageJ. Where possible, three biological replicates were taken for each nematode species with and without *A. oligospora* and multiple technical replicate images were measured within each biological replicate. Statistical analysis was carried out in GraphPad Prism

### Measuring nematode movement

10 adult nematodes were placed in the centre of an NGA plate in 100µl of dd water using a worm pick. The plates were then filmed for approximately 20 seconds with a Nikon D3400 camera and Tamron SP Di AF 90mm macro lens in a dark cabinet with one controlled light source. Three plates were made and filmed for each nematode species. 600 frames (10 seconds at 60 frames/second) were selected from each video and the approximate length of each nematode measured using the straight-line tool in Fiji, ImageJ. The Trackmate plugin with the LAP tracker was then used to identify and track worms over the course of the video. The tracks generated were exported and those that corresponded to a nematode were selected. Parameters expressed in pixels or frames were converted to mm and seconds using the in-frame ruler for distance calibration and the known frame rate of the 60 frames/second. 10 complete worm tracks from across the 3 plates videoed for each species were used to calculate average speed. Straight line speed (mm/sec) was the parameter used to define the speed of each nematode species.

### RNAseq data analysis

Cutadapt (version 1.18) (Martin, 2011) was used to remove adapters off raw reads which were then aligned and sorted into bam files using STAR (Dobin et al., 2013) using reference genomes from *S. ratti* (PRJEB125; WormBase ParaSite), *C. elegans* (PRJNA13758; WormBase ParaSite), *C. inopinata* (PRJDB5687; WormBase ParaSite) or *A. oligospora* (GCA_000225545; EnsemblFungi) and GTF annotations. Reads were counted using HTSeq-count (Anders et al., 2015).

### Differential Expression analysis

Differentially expressed reads were identified using DESeq2 (Love et al., 2014) and implemented in R (version 4.2.1) between nematode vs nematode in presence of fungi and fungi vs fungi in presence of nematodes. Only reads with at least 10 reads in at least two replicates were kept and normalised by DESeq2’s median of ratios (Anders and Huber, 2010). Genes were considered differentially expressed if the p adjusted value < 0.01 and log fold change (FC) > 1.

### Functional annotations

Gene Ontology (GO) terms and gene annotations were predicted using EggNOG-mapper (version 2.1.11) (Cantalapiedra et al., 2021). Functional protein annotations were predicted using InterProScan (version 5) for *S. ratti*, *A. oligospora* and *C. inopinata* and downloaded from WormBase for *C. elegans.* GO terms enriched for significantly upregulated genes were predicted using the TopGO (version 2.40.0) package (Alexa and Rahnenführer, 2020) implemented in R (version 4.2.1). GO terms in Biological Processes (BP), Cellular Components (CC) and Molecular Function (MF) were visualised using a customised REVIGO script (Supek et al., 2011). Venny (2.1.0) used for Venn diagrams.

