## Supplemental Information for "Nematode species differ in susceptibility and response to the nematode-trapping fungus *Arthrobotrys oligospora*"

**Supplementary Information**

*Strongyloides ratti* induces sporulation by the nematode trapping fungus *Arthrobotrys oligospora*

**Key words:**

**Contents Page:**

1. Supplementary Figure 1 – trap formation in response to three dead nematode species analysis of *S. ratti*, *C. inopinata* and *C. elegans*
2. Supplementary Figure 2 – movement analysis of *C. elegans*, *C. inopinata* and *S. ratti* and trap induction in *A. oligospora*.
3. Supplementary Figure 3 – Corelation plots between number of traps formed and speed or total displacement.
4. Supplementary Figure 4 - PCA plots from DESeq2 analysis of gene expression comparisons between the three nematode species and controls.
5. Supplementary Figure 5 – DEseq2 differential expression of genes between nematodes incubated with and without *A. oligospora*
6. Supplementary Figure 6 – Counting fungal growth in response to the three nematode species
7. Supplementary Figure 7 – GO terms related to reproduction upregulated in *C. elegans* and downregulated in *S. ratti* when incubated with *A. oligospora*

**SI Figure 1.**

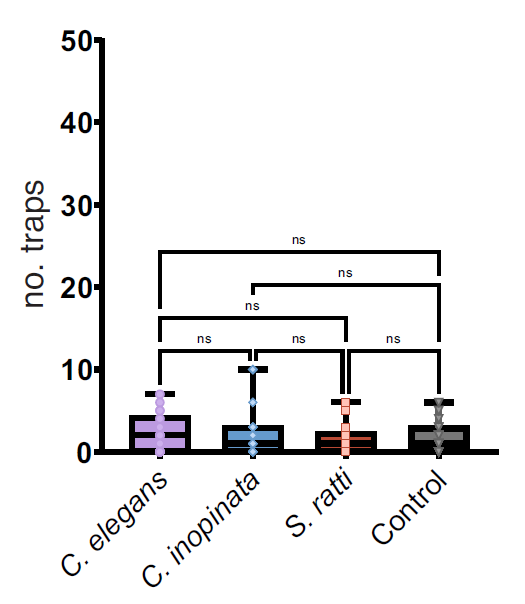

**SI Figure 1 – *A. oligospora* trap formation in response to dead *Caenorhabditis elegans*, *C. inopinata* and *Strongyloides ratti.***

(a) Number of traps formed after 24 hours in response to 10 *C. elegans*, 10 *C. inopinata* or 10 *S. ratti* free-living adult nematodes that have been first killed by freezing.

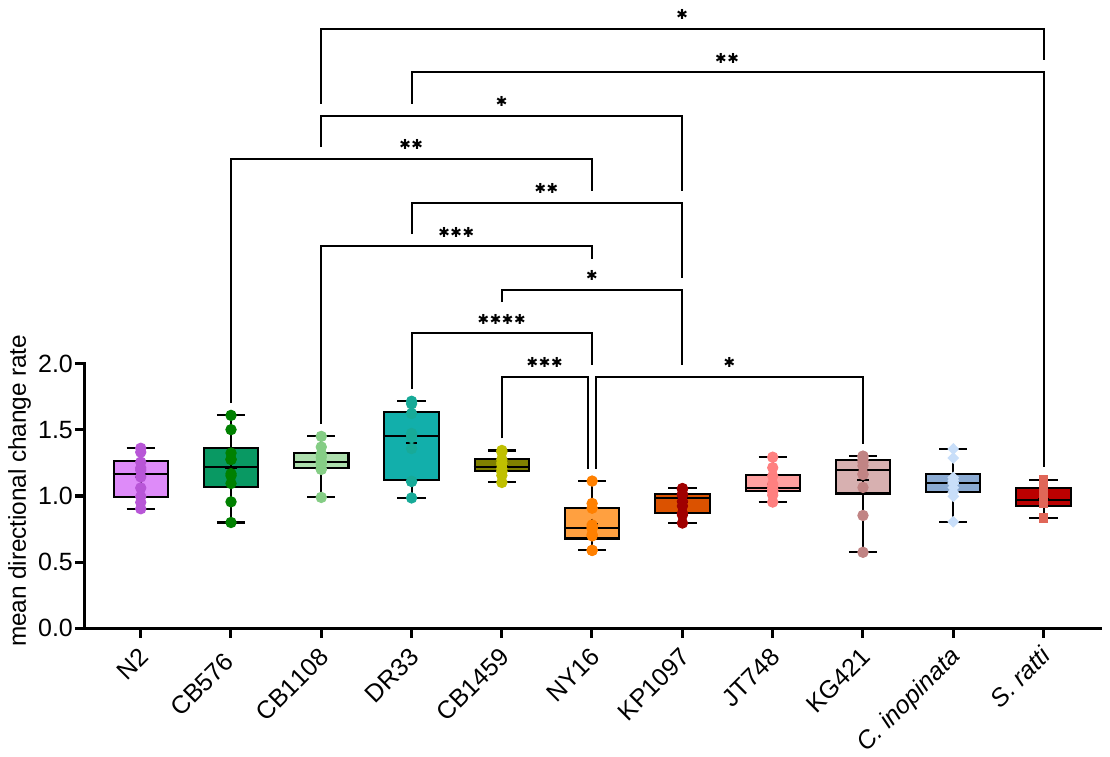

**SI Figure 2 – movement analysis of *C. elegans*, *C. elegans* mutants, *C. inopinata* and *S. ratti***

Directional change have been measured using ImageJ for *C. elegans* N2 lab strain, four hypoactive *C. elegans* (CB576, CB1108, DR33 and CB1459), four hyperactive *C. elegans* lines (NY16, KP1097, JT478 and KG421), *C. inopinata* and *S. ratti.* Statistical analysis calculated with One-way ANOVA with Kruskal-Wallis multiple comparison test.

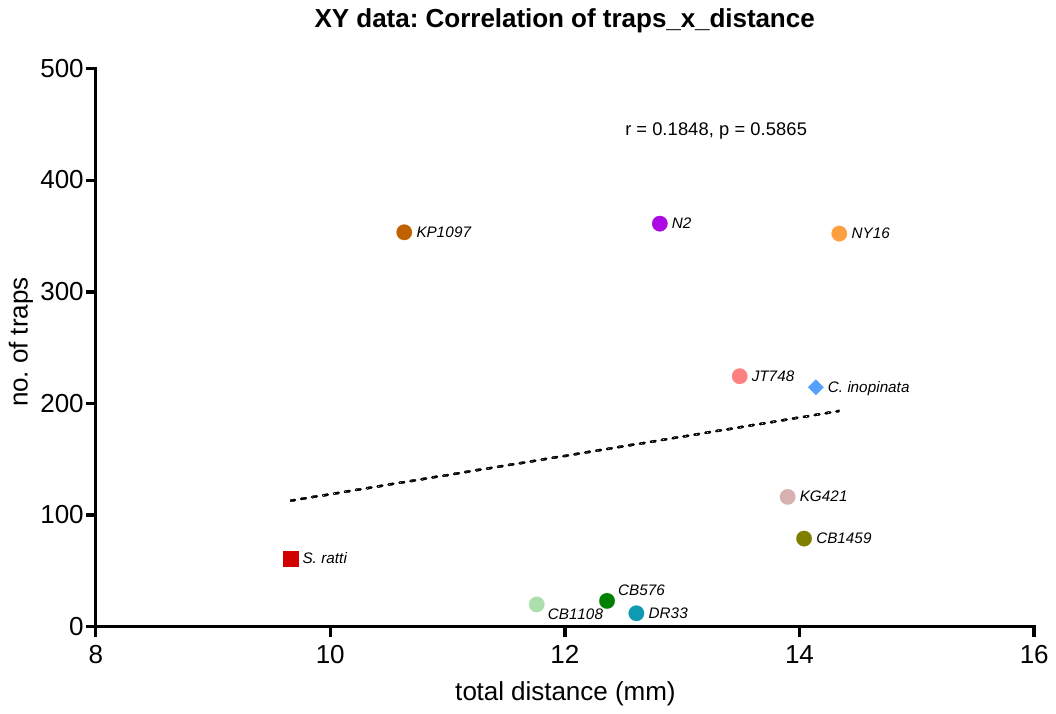

B.

A.

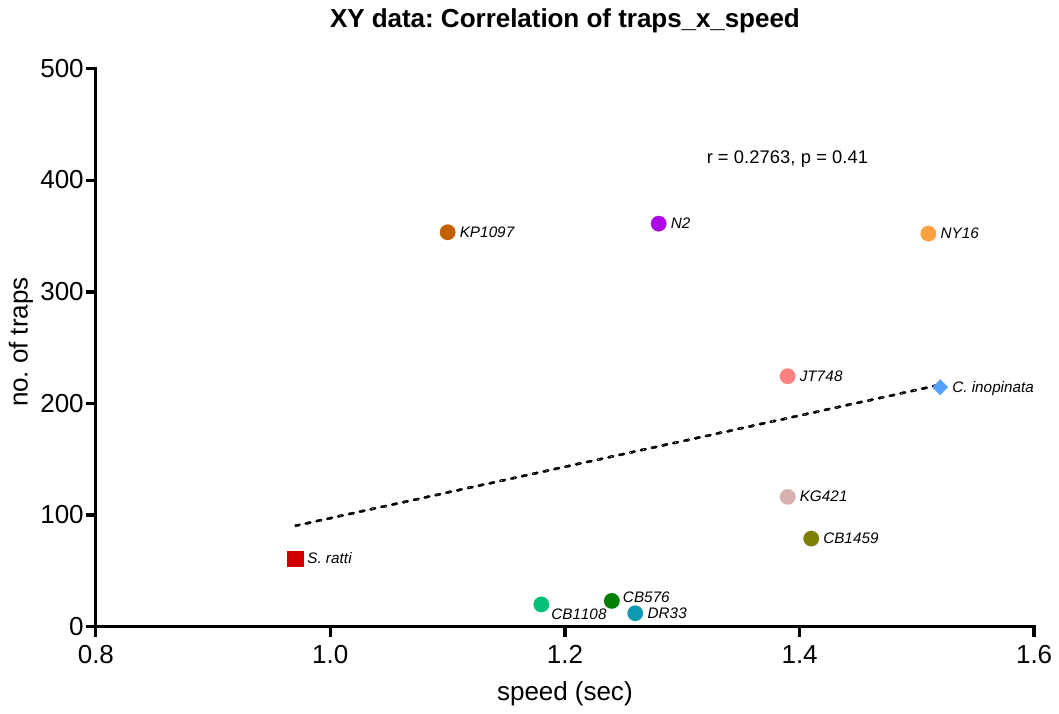

**SI Figure 3 – Correlation between speed, total distance and number of traps**

Pearson’s correlation between number of traps formed by *A. oligospora* and the speed (sec) or total distance (mm) of *S. ratti*, *C. inopinata*, *C. elegans* and the four hypo- and hyper- active *C. elegans* mutants.

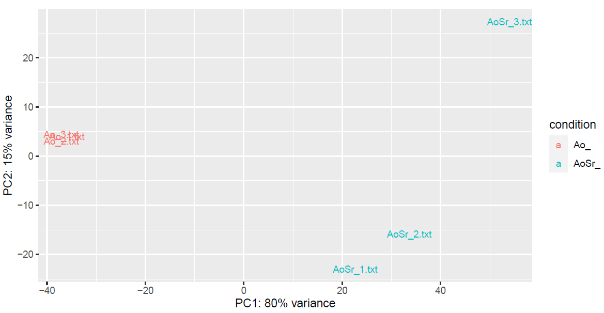

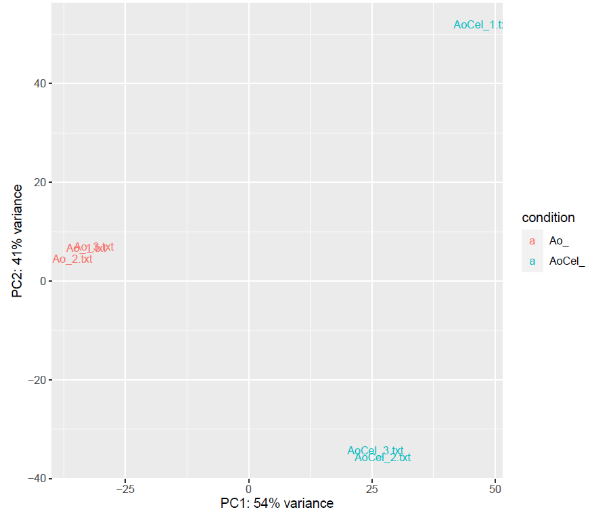

A.

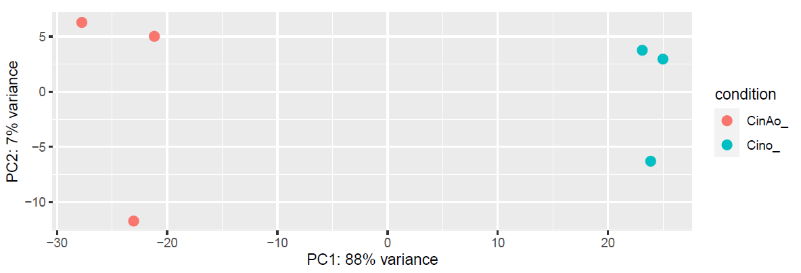

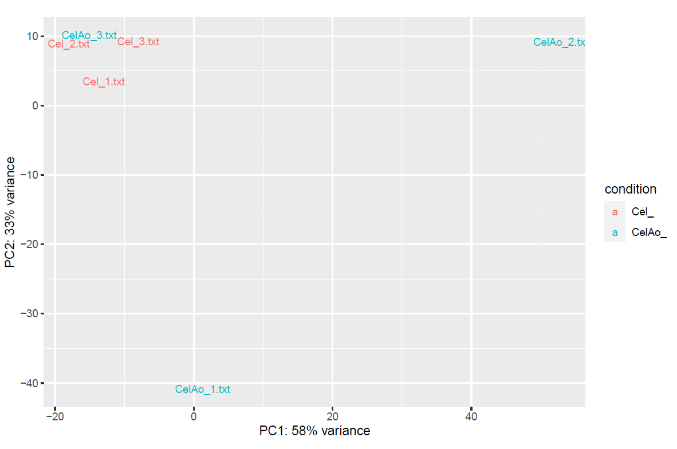

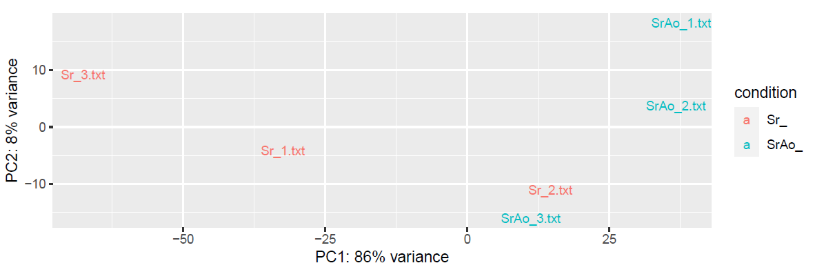

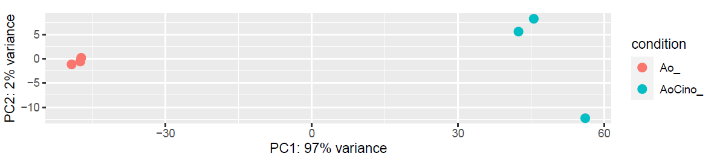

**SI Figure 4 - Correlation and PCA plots from DESeq2 analysis.**

(A) PCA plots of sample divergence for each comparison *A. oligospora* vs *C. elegans,* *S. ratti* and *C. inopinata* shown in A.

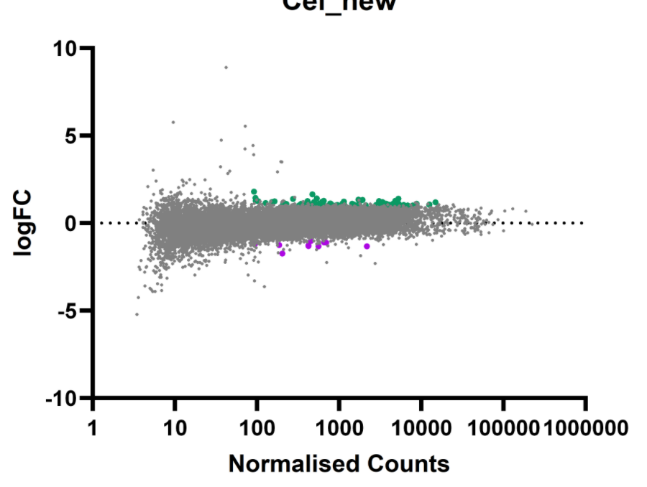

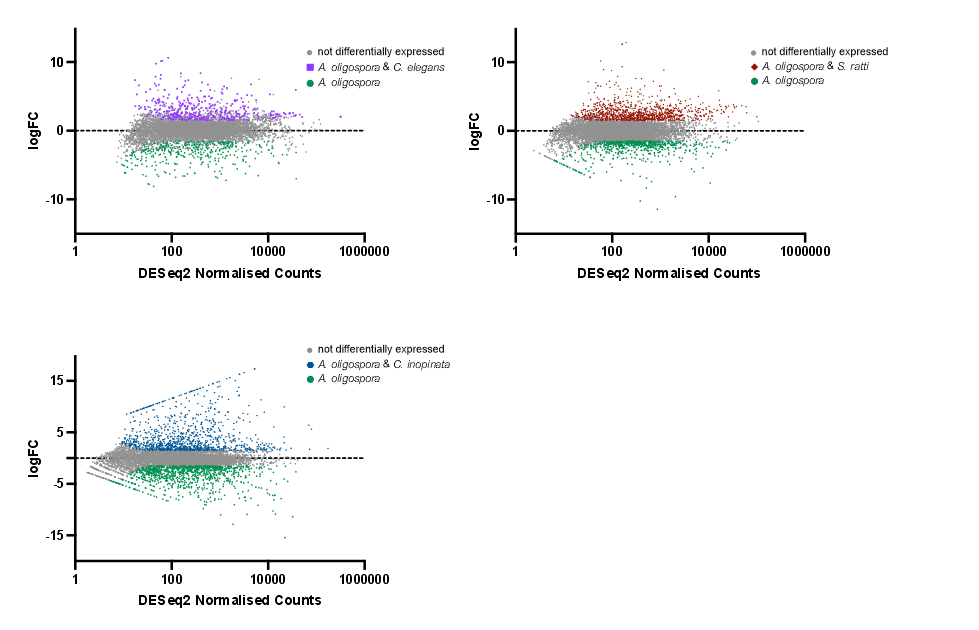

F

E

D

C

B

A

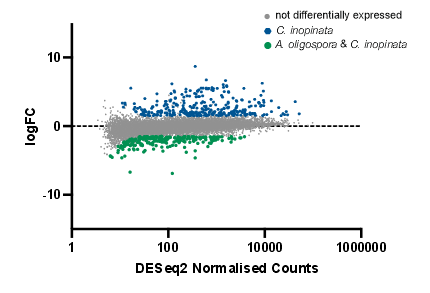

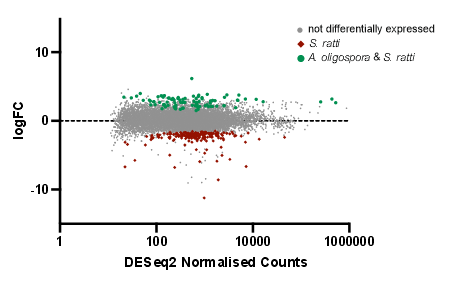

**SI Figure 5 - DESeq2 analysis**

A pairwise comparison of *A. oligospora* (green circles) and *A. oligospora* incubated with either a) *C. elegans* (purple squares), b) *S. ratti* nematodes (red diamonds) or c) *C. inopinata* nematodes (blue circles) and a pairwise comparison of either e) *C. elegans*, f) *S. ratti* or (g) *C. inopinata* incubated with and without *A. oligospora* on axis showing log fold change (log FC) in expression vs normalised gene counts identified using DESeq2 in R. Genes considered significantly upregulated if the adjusted p value ≤ 0.01. Each point is a unique gene.

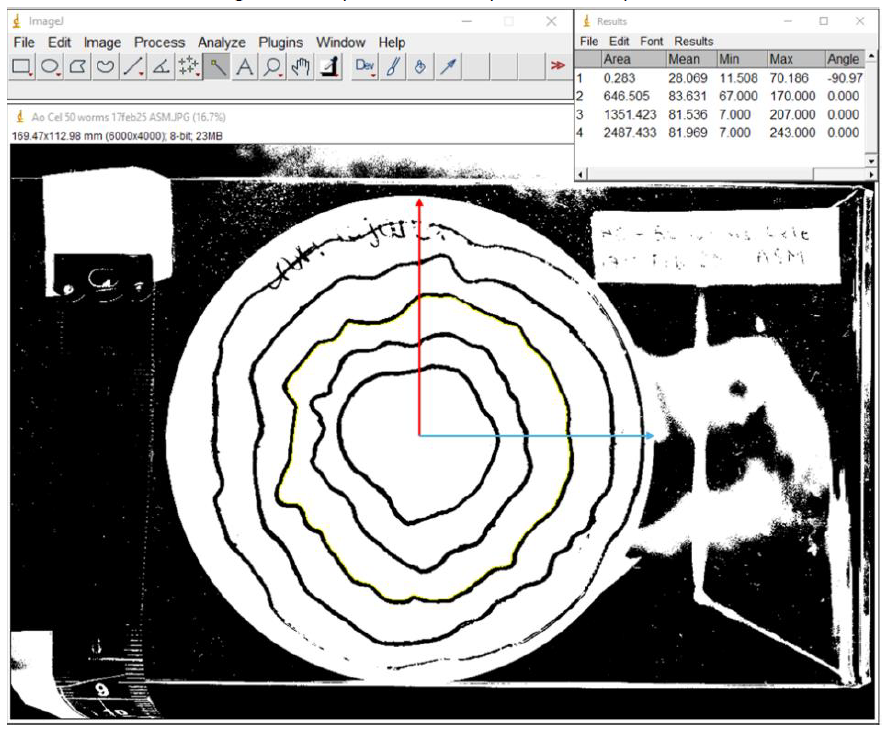

**SI Figure 6 – *A. oligospora* growth rate quantification in response to nematodes**

ImageJ was used to measure radial and area growth of *A. oligospora* on LNA plates in absence and presence of *C. elegans*, *C. inopinata* and *S. ratti* for 15 replicates. The circumference and area of each ring was measured by outward radial measurements (red and blue) for each growth rings are taken in two directions: 0 and 90 degrees. Contour lines (yellow) indicate growth rate.

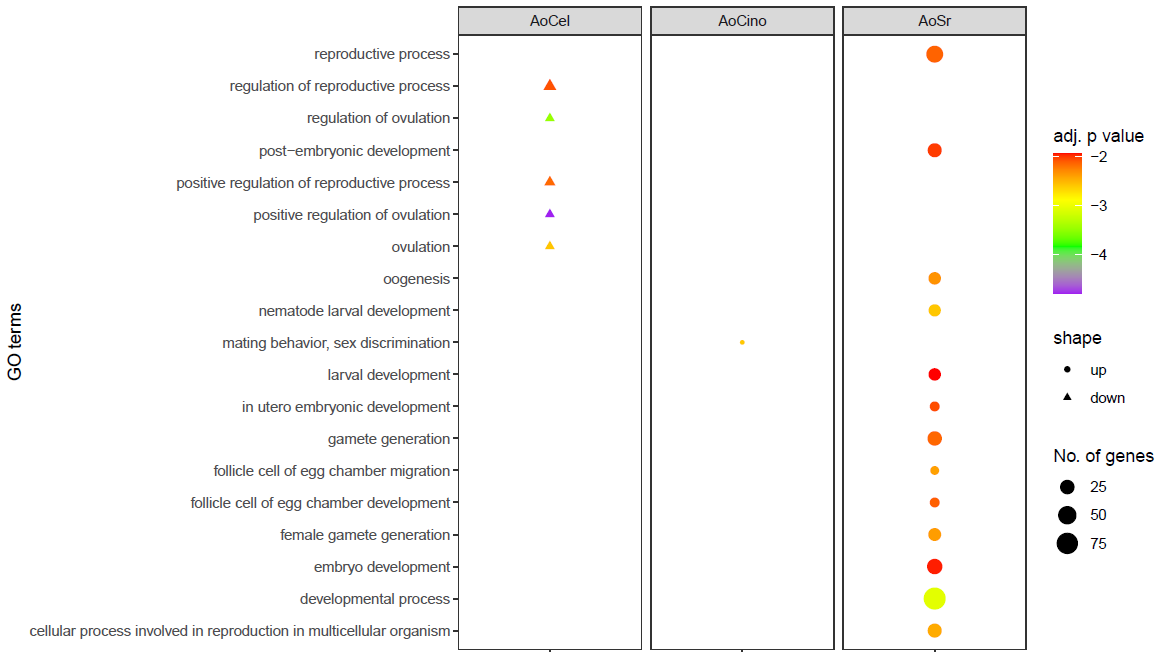

**SI Figure 7 – GO terms related to reproduction upregulated in C. elegans and downregulated in *S. ratti* when incubated with *A. oligospora***

Colour indicated by adjusted p value, size indicated by number of significant genes. Plotted with a custom Revigo script in R.
